# Monovision and the Misperception of Motion

**DOI:** 10.1101/591560

**Authors:** Johannes Burge, Victor Rodriguez-Lopez, Carlos Dorronsoro

## Abstract

Monovision corrections are a common treatment for presbyopia. Each eye is fit with a lens that sharply focuses light from a different distance, causing the image in one eye to be blurrier than the other. Millions of people in the United States and Europe have monovision corrections, but little is known about how differential blur affects motion perception. We investigated by measuring the Pulfrich effect, a stereo-motion phenomenon first reported nearly 100 years ago. When a moving target is viewed with unequal retinal illuminance or contrast in the two eyes, the target appears to be closer or further in depth than it actually is, depending on its frontoparallel direction. The effect occurs because the image with lower illuminance or contrast is processed more slowly. The mismatch in processing speed causes a neural disparity, which results in the illusory motion in depth. What happens with differential blur? Remarkably, differential blur causes a reverse Pulfrich effect, an apparent paradox. Blur reduces contrast and should therefore cause processing delays. But the reverse Pulfrich effect implies that the blurry image is processed more quickly. The paradox is resolved by recognizing that: i) blur reduces the contrast of high-frequency image components more than low-frequency image components, and ii) high spatial frequencies are processed more slowly than low spatial frequencies, all else equal. Thus, this new illusion—the reverse Pulfrich effect—can be explained by known properties of the early visual system. A quantitative analysis shows that the associated misperceptions are large enough to impact public safety.

---

In the year 2020, nearly two billion people in the world will have presbyopia^1^. Presbyopia, a part of the natural aging process, is the loss of focusing ability due to the stiffening of the crystalline lens inside the eye^2^. All people develop presbyopia with age, so the number of affected people increases as the population ages. Without correction, presbyopia prevents people from reading and from effectively using a smartphone.

Many corrections exist for presbyopia. Reading glasses, bifocals, and progressive lenses are well known examples. Less well known are monovision corrections. With a monovision correction, each eye is fitted with a lens that sharply focuses light from a different distance, providing ‘near vision’ in one eye and ‘far vision’ in the other. Monovision thus causes differential blur in the left- and right-eye images of a target at a given distance. For patients in which the correction is successful, the visual system suppresses the lower quality image and preferentially processes the higher quality image^3–5^. The consequence is an increase in the effective depth of field without many of the drawbacks of other corrections (e.g. bifocals cause a ‘seam’ in the visual field). Unfortunately, monovision does not come without its own drawbacks. Monovision degrades stereoacuity^6,7^ and contrast sensitivity^8^, deficits that hamper fine-scale depth discrimination and reading in low light. Monovision is also thought to cause difficulties in driving^9,10^and has been implicated in an aviation accident^11^. Despite these drawbacks, many people prefer monovision corrections to other corrections for presbyopia, or to no corrections at all^12^.

Nearly ten million people in the United States currently have a monovision correction (see Supplement). The number of candidates will increase in the coming years. The population is aging and monovision is the most popular contact lens correction for presbyopia amongst the baby boomers^12^. A full understanding of the effects of monovision on visual perception is critical, both for sound optometric and ophthalmic practice and for the protection of public safety. Unfortunately, there is no literature on how the differential blur induced by monovision impacts motion perception.

We investigated the impact of differential blur on motion perception by measuring the Pulfrich effect, a stereo-motion phenomenon first reported nearly 100 years ago^13^. When a target oscillating horizontally in the frontoparallel plane is viewed with unequal retinal illuminance or contrast in the two eyes, the target appears to move along an elliptical trajectory in depth (Fig. 1A). The effect occurs because the image in the eye with lower retinal illuminance or contrast is processed more slowly than the image in the other eye^13–19^. The mismatch in processing speed causes a neural binocular disparity, a difference in the effective retinal locations of target images in the two eyes^20,21^, which results in the illusory motion in depth.

**Figure 1.**
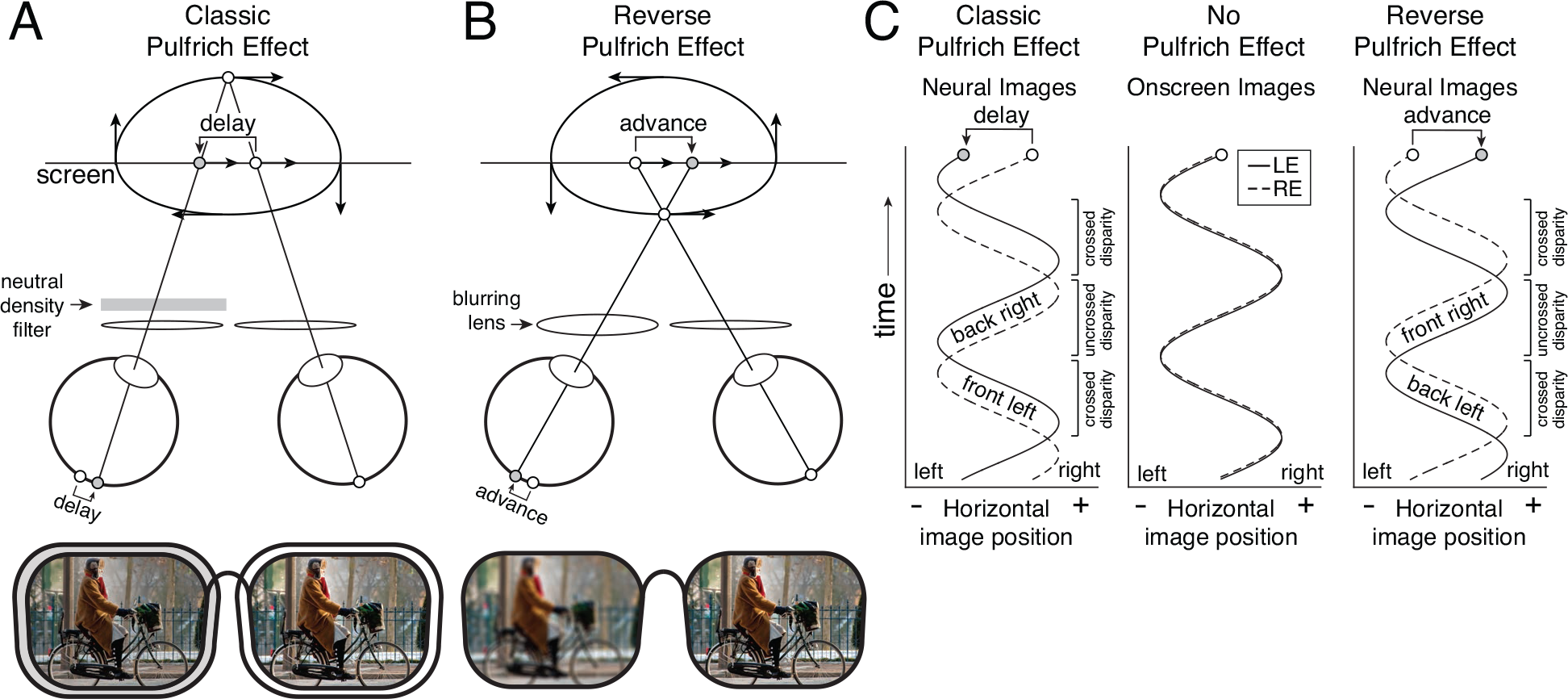
Classic and reverse Pulfrich effects. **A** Classic Pulfrich effect. A neutral density filter in front of the left eye causes sinusoidal motion in the frontoparallel plane to be misperceived in depth (i.e. illusory clockwise motion from above: ‘back right’, ‘front left’). The effect occurs because the response of the eye with lower retinal illuminance (gray dot) is delayed relative to the other eye (white dot), causing a neural disparity. **B** Reverse Pulfrich effect. A blurring lens in front of the left eye causes illusory motion in depth in the other direction (i.e. counter-clockwise from above: ‘front right’, ‘back left’). The effect occurs because the response of the eye with increased blur (gray dot) is advanced relative to the other eye (white dot), causing a neural disparity with the opposite sign. **C** Effective neural image positions in the left and right eye as a function of time for the classic Pulfrich effect, no Pulfrich effect, and the reverse Pulfrich effect.

The Pulfrich effect has been researched extensively since its discovery. The effect is elicited by interocular luminance differences^13^and interocular contrast differences^19^. Its magnitude depends on overall luminance^14,16,22^, dark adaptation^16,23–25^, and numerous other factors^26–28^. In the late 1990s and early 2000s, a flurry of work debated what the effect reveals about the neural basis of stereo and motion encoding^29–33^. But it is not known whether the Pulfrich effect occurs under conditions similar to those induced by monovision corrections.

Do interocular blur differences, like interocular illuminance and contrast differences, cause misperceptions of motion? More specifically, does blur slow the speed of processing and cause a Pulfrich effect? In the classic Pulfrich effect, if the left eye’s retinal illuminance or contrast is decreased, observers perceive ‘front left’ motion (i.e. clockwise motion when viewed from above; Fig. 1A). However, we find that when the left eye is blurred, observers perceive ‘front right’ motion (i.e. counter-clockwise motion when viewed from above; Fig. 1B). Thus, rather than causing a classic Pulfrich effect, differential blur causes a reverse Pulfrich effect.

The discovery of the reverse Pulfrich effect implies an apparent paradox. Blur reduces contrast and should therefore cause the blurry image to be processed more slowly, but the reverse Pulfrich effect implies that the blurry image is processed more quickly than the sharp image (Fig. 1C). At first, this finding appears at odds with a large body of neurophysiological and behavioral results. Low contrast images are known to be processed more slowly at the level of early visual cortex^34–37^ and at the level of behavior^8,38^.

The paradox is resolved by recognizing two facts. First, optical blur reduces the contrast of high spatial frequency image components more than low-frequency image components ^39–42^. Second, extensive neurophysiological^43–45^ and behavioral^8,38^ literatures indicate that high spatial frequencies are processed more slowly than low spatial frequencies, all else equal. Thus, the blurry image is advanced in time relative to the sharp image because the high spatial frequency components in the sharp image decrease the speed at which it is processed. Thus, a new version of a 100-year-old illusion can be explained by known properties of the early visual system.

## Results

To measure the impact of differential blur on the perception of motion we used trial lenses to induce interocular blur differences, and a haploscope for dichoptic presentation of moving targets (Fig. 2A). We measured the strength of the Pulfrich effect using a one-interval two-alternative forced choice (2AFC) experiment. On each trial, a binocular target oscillated sinusoidally from left to right (or right to left) while the observer fixated a central dot. The onscreen interocular delay of the target images was under experimenter control. If the onscreen interocular delay is zero, the stereoscopic target is specified by onscreen disparity to be moving in the plane of the screen. If the onscreen delay is non-zero, it specifies a stereoscopic target moving on an elliptical trajectory outside the plane of the monitor. The task was to report whether the target was moving leftward or rightward when it appeared to be closer than the screen (i.e. clockwise or counter-clockwise when viewed from above; see Fig. 1AB). Human observers made these judgments easily and reliably.

**Figure 2.**
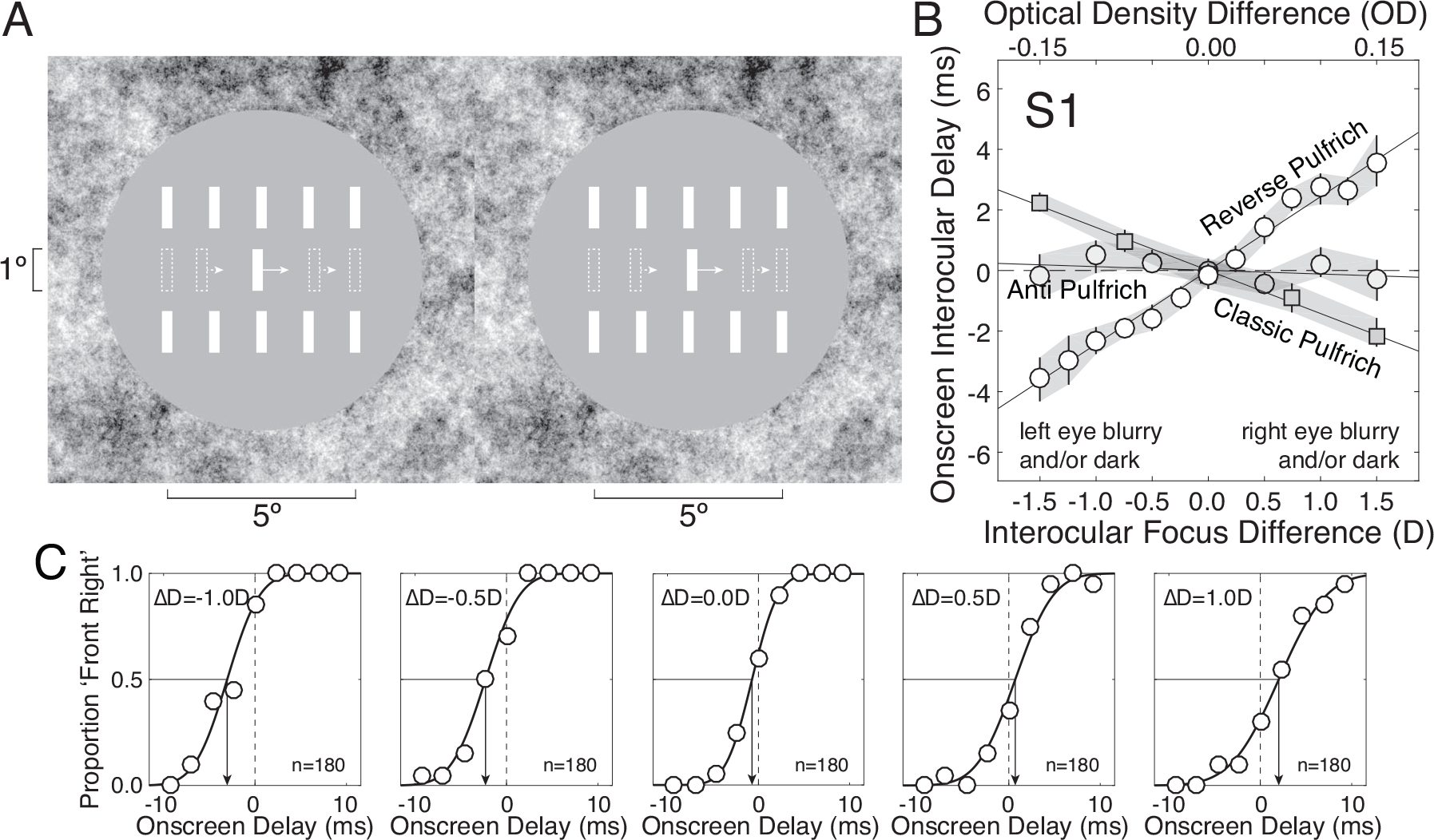
Reverse, classic, and anti-Pulfrich effects. **A** Binocular stimulus. Observers reported whether they saw three-dimensional (3D) target motion as ‘front right’ or ‘front left’ with respect to the screen. Stationary white ‘picket fence’ reference bars served to indicate the screen distance. The target was a horizontally moving 0.25×1.0° white bar. Arrows show motion speed and direction, and dashed bars show bar positions throughout a trial; both are for illustrative purposes only and were not present in the actual stimulus. Fuse the two half-images to perceive the stimulus in 3D. Cross- and divergent-fusers will perceive the bar in front and behind the screen respectively. **B** Points of subjective equality (PSEs) for one human observer, expressed as onscreen interocular delay relative to baseline. Interocular differences in focus error (bottom axis, white circles) cause the reverse Pulfrich effect. Interocular differences in retinal illuminance (top axis, gray squares) cause the classic Pulfrich effect. Appropriately tinting the blurring lens (light gray circles) can eliminate the motion illusions and act as an anti-Pulfrich prescription. (In anti-Puflrich conditions, the optical density was different for each observer at each interocular focus difference.). Shaded regions indicate bootstrapped standard errors. Best-fit regression lines are also shown. **C** Psychometric functions for five of the conditions with differential blur in B. The arrows indicate the raw PSE in each condition.

For a given interocular difference in focus error, we measured the proportion of times that the observers reported ‘front-right’ (i.e. counter-clockwise when viewed from above) as a function of the onscreen interocular delay between the left and right eye images. Performance was summarized with the point of subjective equality (PSE), the 50% point on each psychometric function (Fig. 2BC). The PSE specifies the amount of onscreen interocular delay required to make the target appear to move in the plane of the display screen (i.e. no motion in depth).

The magnitude of the reverse Pulfrich effect increases linearly with the difference in focus error between the two eyes (Fig. 2B, white circles; Fig. S1). Negative interocular differences in focus error indicate conditions in which the left-eye retinal image is blurry and the right-eye retinal image is sharp. In these conditions, the left-eye onscreen image must be delayed (i.e. negative PSE shift) for the target to be perceived as moving in the plane of the screen. Conversely, positive interocular differences in focus error indicate that the left-eye retinal image is sharp and the right-eye retinal image is blurry. In these conditions, the right-eye onscreen image must be delayed (i.e. positive PSE shift). The results indicate that the blurrier image is processed faster than the sharper image. For the first human observer, a ±1.5D difference in focus error caused an interocular difference in processing speed of approximately ±3.7ms (Fig. 2B).

As a control, we measured the classic Pulfrich effect. To do so, we systematically reduced the retinal illuminance to one eye while leaving the other eye unperturbed (see Methods). As expected, the pattern of PSE shifts reverses (Fig. 2B, gray squares; Fig. S1). When the left eye’s retinal illuminance is reduced, the left-eye onscreen image must be advanced in time so that the target is perceived as moving in the plane of the screen, and vice versa. Consistent with classic findings, these results indicate that the darker image is processed more slowly than the brighter image.

Why does the reverse Pulfrich effect occur? To test the hypothesis that the blurry image is processed faster because the high spatial frequencies in the sharp image slow its processing down (see above), we ran an additional experiment with two critical conditions. In the first condition, the onscreen stimulus to one eye was high-pass filtered while the other stimulus was unperturbed. High-pass filtering artificially sharpens the image by removing low frequencies, increases the average spatial frequency, and should decrease the processing speed relative to the original. In the second condition, the onscreen stimulus to one eye was low-pass filtered (Fig. 3AB) while the other was unperturbed. Low-pass filtering removes high frequencies, approximates the effects of optical blur, and should increase the speed of processing relative to the original unperturbed stimulus. Results with high- and low-pass filtered stimuli should therefore resemble the classic and reverse Pulfrich effects, respectively. The predictions are confirmed by the data (Fig. 3C; Fig. S2). Note that these differences in processing speed cannot be attributed to luminance or contrast differences, because the stimuli were designed such that the low-pass filtered stimuli and high-pass filtered stimuli had identical luminance and contrast (Fig. S3). The detailed computational rules that relate frequency content to processing speed remain to be worked out and should make a fruitful area for future study.

**Figure 3.**
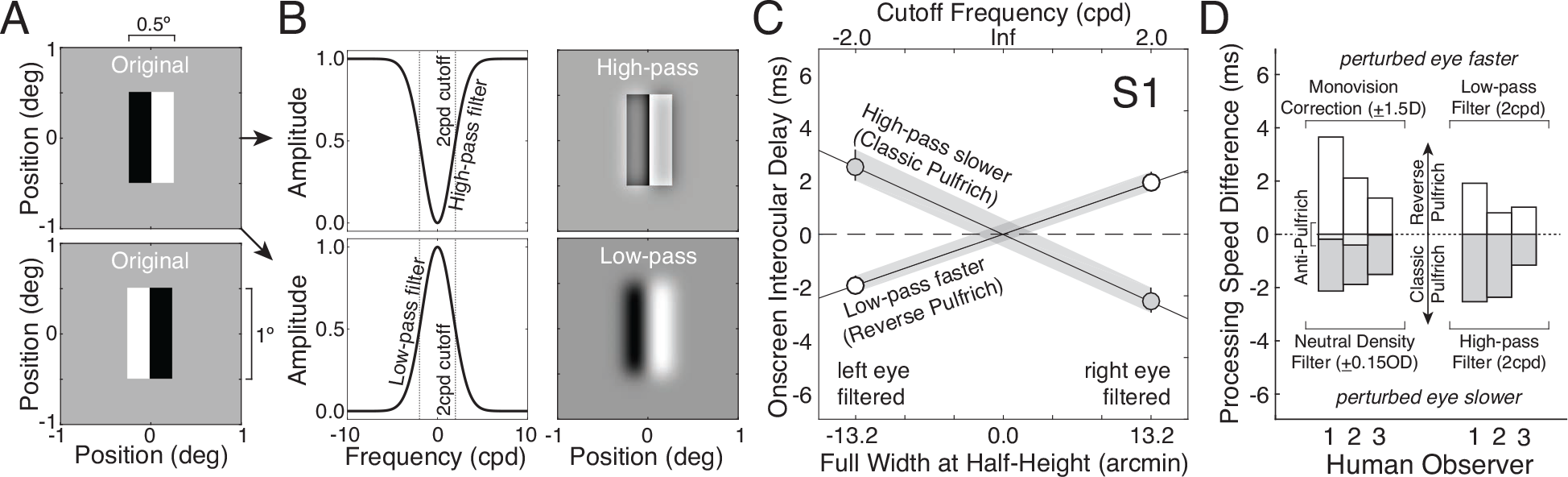
Spatial frequency filtering and the Pulfrich effect. **A** Stimuli to be high- and low-pass filtered were composed of adjacent black-white (top) or white-black (bottom) 0.25°×1.00° bars. **B** High-pass or low-pass filtering changes the spatial frequency content relative to the original (shown only for black-white bar stimuli). High-pass filtered stimuli and low-pass filtered stimuli had identical luminance and contrast (see Fig. S3). **C** High-pass filtered stimuli are processed slower whereas low-pass filtered stimuli are processed faster than the original unperturbed stimulus. **D** Effect sizes for each of three human observers in multiple conditions; effect sizes were obtained from the values of the best-fit regression line in the indicated conditions. Two situations resulted in reverse Pulfrich effects (perturbed image processed faster than unperturbed): interocular focus differences (left, white bars) and low-pass filtering one eye (right, white bars). Two other situations resulted in classic Pulfrich effects (perturbed image processed slower than unperturbed): interocular retinal illuminance differences (left, gray bars) and high-pass filtering one eye (right, gray bars). A fifth situation—appropriately darkening the blurring lens (left, small light gray bars)—eliminates the Pulfrich effect and acts as an anti-Pulfrich correction.

The pattern of performance characterizing the first human observer is consistent across all human observers (Fig. 3D; Figs. S1, S2). The largest differences in focus error (±1.5D) elicited interocular differences in processing speed ranging from 1.4-3.7ms across observers (Fig. 3D, left, white bars). The largest differences in retinal illuminance (±0.15OD) caused differences in processing speed ranging from 1.5-2.1ms across observers. Similar effect sizes are seen for low- and high-pass filtering. These differences in processing speed appear modest. However, a few milliseconds difference in processing speed can lead to dramatic illusions in depth (see below).

The interocular differences in processing speed vary across observers but the magnitude of the differences appear to be correlated across conditions for each observer (Fig. 3D). A larger pool of observers is necessary to confirm whether this trend is significant. Future studies should measure the range and attempt to determine the origin of these inter-observer differences across the human population.

### Motion Illusions in the Real World

Monovision corrections cause misperceptions of motion. How large are these misperceptions likely to be in daily life? If the illusions are small, they will impose no impediment and can be safely ignored. If the illusions are large, they may pose a important issue. To generalize laboratory results to the real world, multiple factors must be taken into account; differences in viewing conditions can change the magnitude of the effects. The same focus error causes less blur with smaller pupils, the same interocular difference in processing speed results in larger binocular disparities at faster speeds, and the same disparity specifies larger depths at longer viewing distances. Thus, all these factors—pupil size, target speed, and viewing distance—must be taken into account when predicting the severity of misperceptions that wearers of monovision corrections are likely to experience in daily life (see Methods).

Consider a target object, five meters away, moving from left to right in daylight conditions. Illusion sizes, as predicted by stereo geometry, are shown (Fig. 4A) for one observer as a function of target speed with different monovision corrections strengths, which typically range between 1.0D and 2.0D^10^. Consider the curve associated with a +1.5D interocular difference in optical power (far lens over left eye), a common monovision correction strength. With this correction, the distance of a target at 5.0 meters moving from left to right at 15 miles per hour will be overestimated by 2.8m (see Methods). This, remarkably, is the width of a narrow street lane! If the prescription is reversed (−1.5D; far lens over right eye), target distance will be underestimated by 1.3m. Stronger monovision corrections and faster speeds will increase the illusion sizes. Illusion sizes are also expected to increase in dim light (e.g. driving at dawn, dusk or night; Fig. S4). This is for two reasons. First, pupil sizes increase in dim light so a given focus error causes more blur^46^. Second, neural differences in processing speed tend to be amplified in dim light^14,25^, so a given blur difference should cause larger illusions in dimmer light.

**Figure 4.**
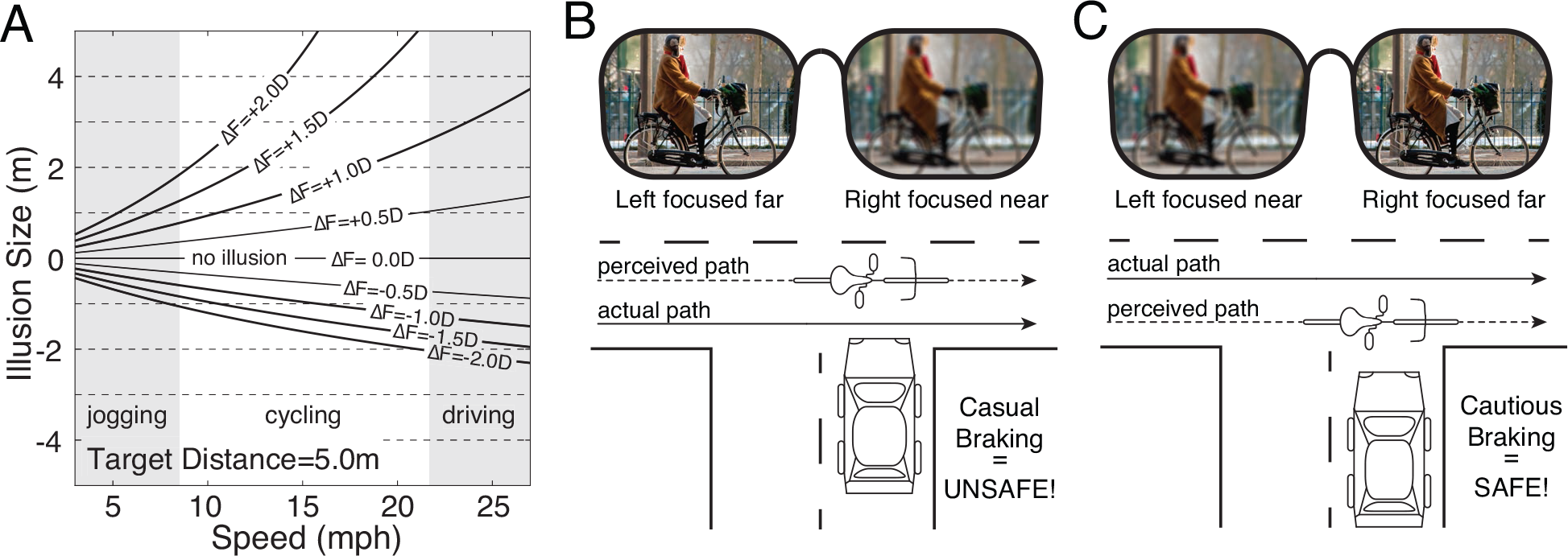
Monovision corrections and misperceptions of depth. **A** Illusion size in meters as a function of speed for an object moving from left to right at 5.0m for different monovision corrections strengths (curves). Monovision correction strengths (interocular focus difference, Δ*F*; see Methods) typically range between 1.0D and 2.0D; strengths of 0.5D are typically not prescribed, but we show them for completeness. Shaded regions show speeds associated with jogging, cycling, and driving. Illusion sizes are predicted directly from stereo-geometry (see Methods) assuming a pupil size (2.1mm) that is typical for daylight conditions^46^, and assuming interocular delays that were measured in the first human observer (see Fig. 2B). The predictions also assume that the observer can sharply focus the target at 5.0m in one eye^2,47^. **B** The distance of cross traffic moving from left to right will be overestimated when the left eye is focused far (i.e. sharp image of cross traffic) and the right eye is focused near (i.e. blurry image of cross traffic). **C** The distance of left to right cross traffic will be underestimated when the left eye is focused near and the right eye is focused far.

Illusions this large will not only be disturbing for the person wearing the monovision correction; they have the potential to compromise public safety. In countries where motorists drive on the right side of the road (e.g. USA), cars and cyclists approaching in the near lane of cross traffic move from left to right. Placing the far lens in the left eye will cause distance overestimation, which may result in casual braking and increase the likelihood of traffic accidents (Fig. 4B). Placing the far lens in the right eye may be advisable. The resulting distance underestimation should result in more cautious braking and reduce the likelihood of collisions (Fig. 4C). In countries where motorists drive on the left side of the road (e.g. United Kingdom), the opposite practice should be considered (i.e. far lens in left eye). The current standard is to place the far lens in the dominant eye^10,48^, but this does not appear to increase patient acceptance rate, patient satisfaction^48,49^, or quantitative measures of visual performance^6,50^. Although the scenarios just discussed are not the only scenarios that ought be considered, they may invite reexamination of standard optometric practice.

In the real world, many monocular cues exist that tend to indicate the correct rather than illusory depths. The literature on cue combination^51,52^ suggests the magnitude of the reverse Pulfrich effect may be somewhat reduced from the predictions in Fig. 4A, but they should still serve as a useful benchmark for comparison. It will be of clinical and scientific interest to examine how the reverse Pulfrich effect manifests in the rich visual environment of the real world^9^. This question could be examined with virtual- or augmented-reality headsets that are capable of providing the experimenter precise programmatic control of near-photorealistic graphical renderings.

Another implication of these results is that objects moving towards an observer along a straight line should appear to follow S-curve trajectories (Fig. S5). These misperceptions should make it difficult to play tennis, baseball, and other ball sports requiring accurate perception of rapidly moving targets. Monovision corrections should be avoided when playing these sports.

## Discussion

### Eliminating monovision-induced motion illusions

Reconsidering prescribing practices is one approach to minimizing the negative consequences of monovision-induced motion illusions, but it is clearly not the perfect solution. It would be far better to eliminate the illusions altogether. Because increased blur and reduced retinal illuminance have opposite effects on processing speed, it should be possible to null the two effects by tinting the lens that forms the blurry image. We reran the original experiment, with appropriately tinted lenses for each human observer (see Methods). This ‘anti-Pulfrich prescription’ eliminates the motion illusion in all human observers (Figs. 2B, 3D).

For any given monovision prescription, the lens forming the blurry image varies according to target distance. Anti-Pulfrich prescriptions thus cannot work for all target distances. Tinting the near lens (blurry, dark images for far targets; sharp, dark images for near targets) will eliminate the Pulfrich effect for far targets but exacerbate it for near targets. However, because many presbyopes have some residual accommodation and because they tend to use it to focus the distance-corrected eye^2,47,53^, the range of far distances over which motion misperceptions can be eliminated may be quite large: 0.67m to the horizon for a patient with 1.5D of residual accommodation. Given that accurate perception of moving targets is likely to be more critical for tasks at far distances (e.g. driving) than at near distances (e.g. reading), tinting the near lens is likely to be the preferred solution. This issue, however, clearly needs further study.

### Adaptation

Previous studies have shown that blur perception changes with consistent exposure to blur^54^. Do motion illusions change as patients adapt to monovision corrections over time? The question has never been asked in the context of the reverse Pulfrich effect. The literature on the classic Pulfrich effect is mixed. At short time scales (i.e. minutes), motion illusions remain unchanged^16,55,56^ or increase^24^ until reaching an asymptote at steady state. At long time scales (i.e. days), motion illusions decrease as observers adapt to interocular differences in light level^57,58^. Also, in these previous adaptation studies, the eye with the dark image was always the same. With a monovision correction, the eye with the blurry image varies with target distance. Thus, it is unclear whether observers will adapt such that motion illusions caused by the reverse Pulfrich effect will be reduced. This is an important area for future study, both for basic science and for the development of successful clinical interventions.

### Spatial frequency binding problem

Like many scientific discoveries, the reverse Pulfrich effect presents new scientific opportunities. We have demonstrated that the reverse Pulfrich effect occurs because sharp images contain more high frequencies (i.e. fine details) than blurry images, and because high frequencies are processed more slowly than low frequencies. Thus, the different frequency components in one eye’s image of a moving target should arrive in cortex at different times. Without a temporal alignment mechanism, the different spatial frequency components in the image should appear to split apart when a target object moves. However, this percept is not typically experienced. The visual system must therefore bind together the different frequency components of a moving stimulus to achieve a unified percept.

Variants of the psychophysical paradigm that we have used to investigate the reverse Pulfrich effect have great potential for investigating the visual system’s solution to the spatial frequency-binding problem. The measurements have exquisite temporal precision, often to within a small fraction of a millisecond (see Fig. 2BC, Fig. 3CD), and this precision should prove useful for studying this important but understudied problem in vision and visual neuroscience.

## Conclusion

We have reported a new version of a 100-year-old illusion: the reverse Pulfrich effect. We found that interocular differences in image blur, like those caused by monovision corrections, cause millisecond interocular differences in processing speed. For moving targets, these differences can cause dramatic illusions of motion in depth. The fact that a mismatch of a few milliseconds can yield substantial misperceptions highlights how exquisitely the visual system must be calibrated for accurate percepts to occur. The fact that these motion illusions are rare indicates how well the visual system is calibrated under normal circumstances.

## Author Contributions

JB wrote the paper. JB and VRL collected and analyzed data. JB, VRL, and CD conceived the project and edited the paper.

## Acknowledgments

We thank Bill Geisler, David Brainard, Josh Gold, Marty Banks, and Mike Landy for comments on a draft version of the manuscript. We thank Larry Cormack for assistance in the construction of the haploscope and VPixx technical support for writing custom firmware to support simultaneous presentation of high-resolution images on two monitors at high frame rates. We thank Benjamin Chin and Takahiro Doi for assistance calibrating the monitors. This work was supported by US NIH grant R01-EY028571 to JB from the National Eye Institute and the Office of Social and Behavioral Science, startup funds to JB from the University of Pennsylvania, Spanish Government Grants FPU17/02760 to VRL from the Ministry of Education, and ISCIII DTS2016/00127 and FIS2017-84753-R to CD from the Ministry of Science, Innovation and Universities.

## Methods

### Observers

Three human observers ran in the experiment; two were authors. All human observers had normal or corrected to normal visual acuity (20/20), a history of isometropia, and normal stereoacuity as confirmed by the Titmus Stereo Test.

### Apparatus

Stimuli were displayed on a custom-built four-mirror haploscope. Left- and right-eye images were presented on two identical VPixx VIEWPixx LED monitors. Monitors were calibrated (i.e. the gamma functions were linearized) using custom software routines. The monitors had a size of 52.2×29.1cm, spatial resolution of 1920×1080 pixels, a native refresh rate of 120Hz, and a maximum luminance of 105.9 cd/m^2^. The maximum luminance after light loss due to mirror reflections was 93.9 cd/m^2^. The monitors were daisy-chained together and controlled by the same AMD FirePro D500 graphics card with 3GB GDDR5 VRAM, to ensure that the left and right eye images were presented synchronously. Custom firmware was written so that each monitor was driven by a single color channel; the red channel drove the left monitor and the green channel drove the right monitor. The single-channel drive to each monitor was then split to all three channels to enable gray scale presentation. Simultaneous measurements with two optical fibers connected to an oscilloscope confirmed that the left and right eye monitor refreshes occurred within ~5 microseconds of one another.

Human observers viewed the monitors through mirror cubes with 2.5cm circular openings positioned one inter-ocular distance apart. Heads were stabilized with a chin and forehead rest. The haploscope mirrors were adjusted such that the vergence distance matched the distance of the monitors. The light path from monitor to eye was 100cm, as confirmed both by a laser ruler measurement and by a visual comparison with a real target at 100cm. At the 100cm viewing distance, each pixel subtended 1.09 arcmin. Anti-aliasing enabled sub-pixel resolution that permitted accurate presentations of disparities as small as 15-20arcsec.

### Stimuli

The target stimulus was a binocularly presented, horizontally moving, white vertical bar (Fig. 2A). The target bar subtended 0.25×1.00° of visual angle. In each eye, the image of the bar moved left and right with a sinusoidal profile. An interocular phase shift between the left- and right-eye images introduced a spatial disparity between the left- and right-eye bars. The left- and right-eye onscreen bar positions were given by

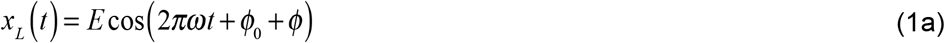

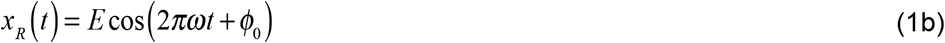

where *x*_*L*_ and *x*_*R*_ are the left and right eye x-positions in degrees of visual angle, *E* is the movement amplitude in degrees of visual angle, *ω* is the temporal frequency, *ϕ*_0_ is the starting phase which in our experiment determines whether the target starts on the left or the right side of the display, *t* is time, and *ϕ* is the phase shift between the images.

The interocular temporal shift (i.e. delay or advance) in seconds associated with a particular phase shift is

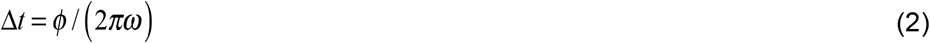

Negative values indicate the left eye onscreen image is delayed relative to the right; positive values indicate the left eye onscreen image is advanced relative to the right.

When the interocular temporal shift equals zero, the virtual bar moves in the fronto-parallel plane at the distance of the monitors. When the interocular temporal shift is non-zero, a spatial binocular disparity results, and the virtual bar follows a near-elliptical trajectory of motion in depth. The binocular disparity in radians of visual angle as a function of time is given by

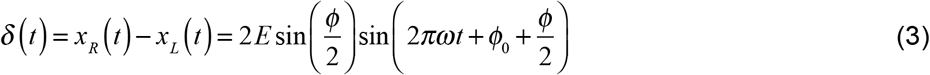

Here, negative disparities are crossed and positive disparities are uncrossed, indicating that the target is nearer and farther than the screen distance, respectively. The binocular disparity takes on its maximum magnitude when the perceived stimulus is directly in front of the observer and the lateral movement is at its maximum speed. When the stimulus is moving to the right, the maximum disparity in visual angle is given by *δ*_max_ = 2*E*sin(*ϕ*/2).

In our experiment, the movement amplitude was 2.5° of visual angle (i.e. 5.0° total change in visual angle in each direction), the temporal frequency was 1 cycle per second, and the starting phase *ϕ*_0_ was randomly chosen to be either 0 or π. Restricting the starting phase to these two values forced the stimuli to start either 2.5° to the right or 2.5° to the left of center on each trial. The onscreen interocular phase shift ranged between +216 arcmin at maximum, corresponding to interocular delays of +10.0 ms. The range and particular values were adjusted to the sensitivity of each human observer.

Two sets of five vertical bars (each identical to the target bar) in a ‘picket fence’ arrangement vertically flanked the region of the screen that was traversed by the target bar. The picket fences were defined by disparity to be at screen distance, and served as a stereoscopic reference for the observer. A 1/f noise texture, also defined by disparity to be at the screen distance, covered the periphery of the display to aid binocular fusion. A small fixation dot marked the center of the screen.

### Procedure

The observer’s task was to report whether the target bar appeared to move leftward or rightward when the stimulus was nearer than the screen in its virtual trajectory in depth. Observers fixated a small centrally located white dot while performing the task. Using a one-interval two-alternative forced choice procedure, nine-level psychometric functions were collected in each condition using the method of constant stimuli. Each function was fit with a cumulative Gaussian using maximum likelihood methods. The 50% point on the psychometric function—the point of subjective equality (PSE)—indicates the onscreen interocular delay needed to null the interocular difference in processing speed. The pattern of PSEs across conditions was fit via linear regression, yielding a slope and y-intercept. Average y-intercepts were nearly zero for each observer: 0.06ms, −0.06ms, and 0.01ms, respectively. To emphasize the differences in slope (i.e. the changes in processing speed in the slope) induced by interocular perturbations, we zeroed the y-intercepts when plotting the PSE data. Observers responded to 180 trials per condition in counter-balanced blocks of 90 trials each.

### Defocus and blur

The interocular focus difference is the magnitude of the defocus in the right eye minus the magnitude of the defocus in the left eye

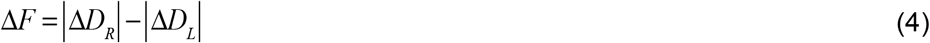

where Δ*D* = *D*_*focus*_ − *D*_*target*_ is the defocus, the difference between the dioptric distances of the focus and target points. To manipulate the amount of defocus blur in each eye, we positioned trial lenses ~12mm from each eye, centered on each optical axis, between each eye and the front of the mirror cubes of the haploscope.

Human observers ran in thirteen conditions defined by interocular focus difference. (One observer, S2, ran in only seven). Each eye was myopically defocused from 0.00D to 1.50D in 0.25D steps while the other eye was kept sharp. Six conditions introduced focus error in the left eye (0.25D to 1.50D in 0.25D steps) while the right eye was sharp (Δ*F* <0.0D). In one condition, both eyes were sharp (Δ*F* =0.0D). And six other conditions introduced focus error in the right eye (0.25D to 1.50D in 0.25D steps) while the left eye was sharp (Δ*F* >0.0D).

In the condition in which both eyes were sharply focused, the optical distances of the left- and right-eye monitors were set to optical infinity with +1.00D trial lenses. When human observers fully relaxed the accommodative power of their eyes, the monitor was clearly focused. All observers indicated that they could sharply focus the monitor under these conditions. Also, because each trial lens absorbs a small fraction of the incident light, having a trial lens in front of each eye in all conditions ensures that retinal illuminance is matched in both eyes in all conditions. To induce interocular differences in focus error, we positioned a stronger positive lens (i.e. +1.25D, +1.50D, +1.75D, +2.00D) in front of one eye. This procedure places one eye’s monitor beyond optical infinity, thereby introducing myopic focus errors that cannot be cleared by accommodation. Before each run, the observer viewed a test target to confirm that he/she could clearly focus targets at optical infinity in the 0.0D baseline condition. Undercorrected hyperopia or overcorrected myopia could place the far point of each eye beyond optical infinity, thereby frustrating our attempts to control the optical conditions. To protect against this possibility, before running each observer, we estimated the far points of the eyes with standard optometric techniques. Then, if necessary, we adjusted the trial lens power so that the monitors were positioned at the desired optical distance.

Another potential concern is that the eyes could accommodate independently to clear the blur in each eye. There are several reasons to think this was not in fact an issue in the current experiments. First, accommodation in the two eyes tends to be strongly coupled, especially for targets straight ahead at distances beyond 1.0m^2,53,59–62^. Second, positioning the optical distance of one monitor beyond optical infinity (see above) minimizes the possibility that differential optical power could be compensated by differential accommodation. Third, discrimination thresholds (*d′* = 1.0) increase systematically with interocular difference in focus error, which is consistent with the literature showing that differential blur deteriorates stereoacuity^63^(see Fig. S6). Fourth, the consistent pattern of results across conditions and observers suggest differential blur was successfully induced.

### Neutral Density Filters

To induce interocular differences in retinal illuminance we placed ‘virtual’ neutral density filters in front of the eyes. To do so, we converted optical density to transmittance, the proportion of incident light that is passed through the filter, using the standard expression *T* = 10^−*OD*^ where *T* is the transmittance and *OD* is the optical density. Then, we reduced the luminance of one eye’s monitor by a scale factor equal to the transmittance. In all observers, performance with real and equivalent virtual neutral density filters is essentially identical, suggesting that the virtual filters were implemented accurately.

The interocular difference in optical density Δ*O* = *OD*_*R*_ −*OD*_*L*_ is the difference between the optical density of filters placed over the right and left eyes. Human observers ran in five conditions with virtual neutral density filters, with equally spaced interocular differences in optical density between −0.15 and 0.15. Two conditions introduced a filter in front of the left eye (Δ*O* <0.00). In one condition, both eyes were unfiltered (Δ*O* = 0.00). And two other conditions introduced a filter in front of the right eye (Δ*O* >0.00)

### Low- and high-pass spatial filtering

To test the hypothesis that the reverse Pulfrich effect is caused by differences in the processing speed of different spatial frequencies, we filtered the onscreen stimulus of one eye with two different frequency filters. The low-pass filter was Gaussian-shaped

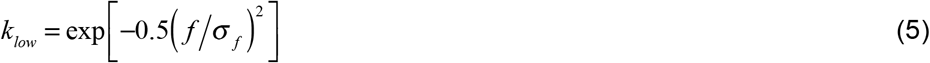

with a standard deviation 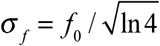 set by the cutoff frequency *f*_0_ such that the filter reached half-height at 2cpd. The high-pass filter complemented the low-pass filter and was given by

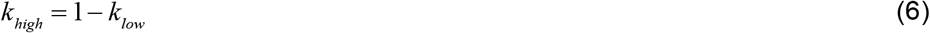

After high-pass filtering the mean luminance was added back in so that the high-pass and low-pass filtered stimuli had the same mean luminance.

To isolate the impact of spatial frequency content on processing speed, it was necessary to modify the onscreen stimulus from the main experiment. Rather than the 0.25°×1.00° white bar, the onscreen stimulus was changed to a 0.50°×1.00° stimulus that was composed of adjacent 0.25×1.0° black and white (or white and black) bars (Fig. 3A). This modification ensured that the low- and high-pass filtered stimuli had identical luminance and identical contrast (see Fig. S3). Each human observer collected 180 trials in each of eight conditions—low- vs. high-pass filtering, left- vs. right-eye filtered, black-white vs. white-black stimulus types—collected in counter-balanced order. Black-white vs. white-black stimulus types had little impact so results were collapsed across stimulus type.

### Generalizing laboratory results to the real world

To make predictions for how monovision corrections will cause misperception of motion in the real world, it is important to take into account the differences in viewing conditions that may impact the magnitude of the effects. The conditions were chosen based on interocular differences in focus error, but the reverse Pulfrich effect is more directly mediated by differences in image blur. The amount of retinal image blur in each eye depends both on the defocus and on the pupil diameter. Thus, it is important to account for changes in pupil diameter due to luminance differences between the lab and the viewing conditions of interest.

The diameter of the blur circle in radians of visual angle is given by

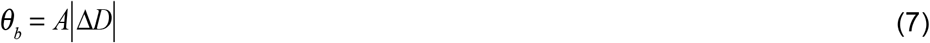

where *θ*_*b*_ is the diameter of the blur circle in radians of visual angle, *A* is the pupil aperture (diameter) in meters and Δ*D* =*D*_*focus*_ − *D*_*target*_ is the defocus which is given by the difference between the dioptric distances of the focus and target points (Fig. S7). In our experiments, we assumed a pupil diameter of 2.5mm, corresponding to the luminance in the experiment^46^. Under the geometrical optics approximation, the absolute value of the defocus |Δ*D*| in the blurry eye equals the absolute value of the interocular focus difference |Δ*F*| because one eye was always sharply focused (i.e. min(|Δ*D*_*L*_, |Δ*D*_*R*_| = 0.0*D*) in our experiments.

The interocular delay in seconds is linearly related to each level of blur by

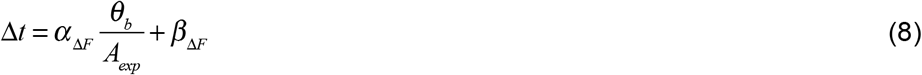

where *α*_*ΔF*_ and *β*_*ΔF*_ are the slope and constant of the best-fit line to the data in Fig. 2B, and *A*_*exp*_ is the pupil diameter of the observer in meters during the experiment. The constant (i.e. y-intercept) can be dropped assuming it reflects response bias and not sensory-perceptual bias.

For a target moving at a given velocity in meters per second, a particular interocular difference in processing speed will yield an effective interocular spatial offset (i.e. position difference)

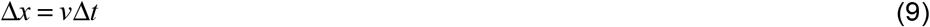

The illusory distance of the target, predicted by stereo-geometry, is given by

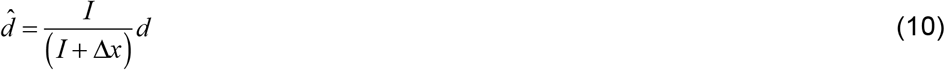

where *d* is the actual distance of the target (Fig. S8).

Combining Eqs. 7–10 yields a single expression for the illusory distance

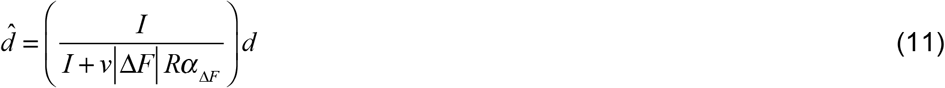

where *R* = *A* / *A*_*exp*_ is the ratio between the pupil diameters in the viewing condition of interest and in the lab when the psychophysical data was collected. Finally, taking the difference between the illusory and actual target distances 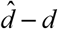 yields the illusion size (see Fig. 4A).

The expression for the illusory distance can also be derived by first computing the neural binocular disparity caused by the delay-induced position difference, and then converting the disparity into an estimate of depth. The binocular disparity in radians of visual angle is given by

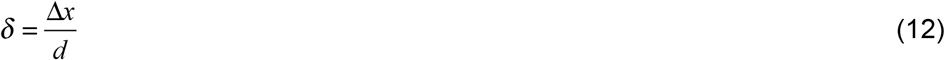

The relationship between illusory distance, binocular disparity, and actual distance is given by

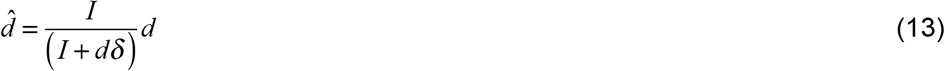

Plugging Eq. 12 into Eq. 13 yields Eq. 10. Thus, both methods of computing the illusory distance are equivalent.

### Anti-Pulfrich monovision corrections

Reducing the image quality of one eye with blur increases the processing speed relative to the other eye and causes the reverse Pulfrich effect. Reducing the retinal illuminance of one eye reduces the processing speed relative to the other eye and causes the classic Pulfrich effect. Thus, in principle, it should be possible to null the two effects by reducing the retinal illuminance of the blurry eye. The interocular delay in seconds is linearly related to each interocular difference in optical density Δ*O* by

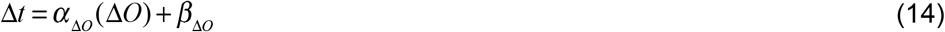

The optical density that should null the interocular delay of a given interocular focus difference is given by

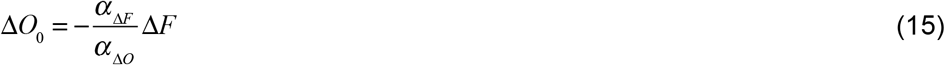

the interocular difference in focus error scaled by the ratio of the slopes of the best-fit linear regression lines to the reverse and classic Pulfrich datasets. The experimental data shows that the optical density predicted by the two regression slopes eliminates the Pulfrich effect (Fig. 2B).

## Supplement

### Prevalence of Monovision Corrections

There are approximately 123 million presbyopes in the USA^64^. Approximately 12.9 million of these presbyopes wear contact lenses, and 4.5 million (35%) of these presbyopic contact lens users have monovision corrections^65,66^. Approximately 30 million presbyopes have had surgery to implant intraocular lenses^67^, and approximately 5.1 million (17%) of these surgical patients have received monovision corrections^68^. Together, this results in approximately 9.6 million presbyopes with monovision corrections in the USA.

**Figure S1.**
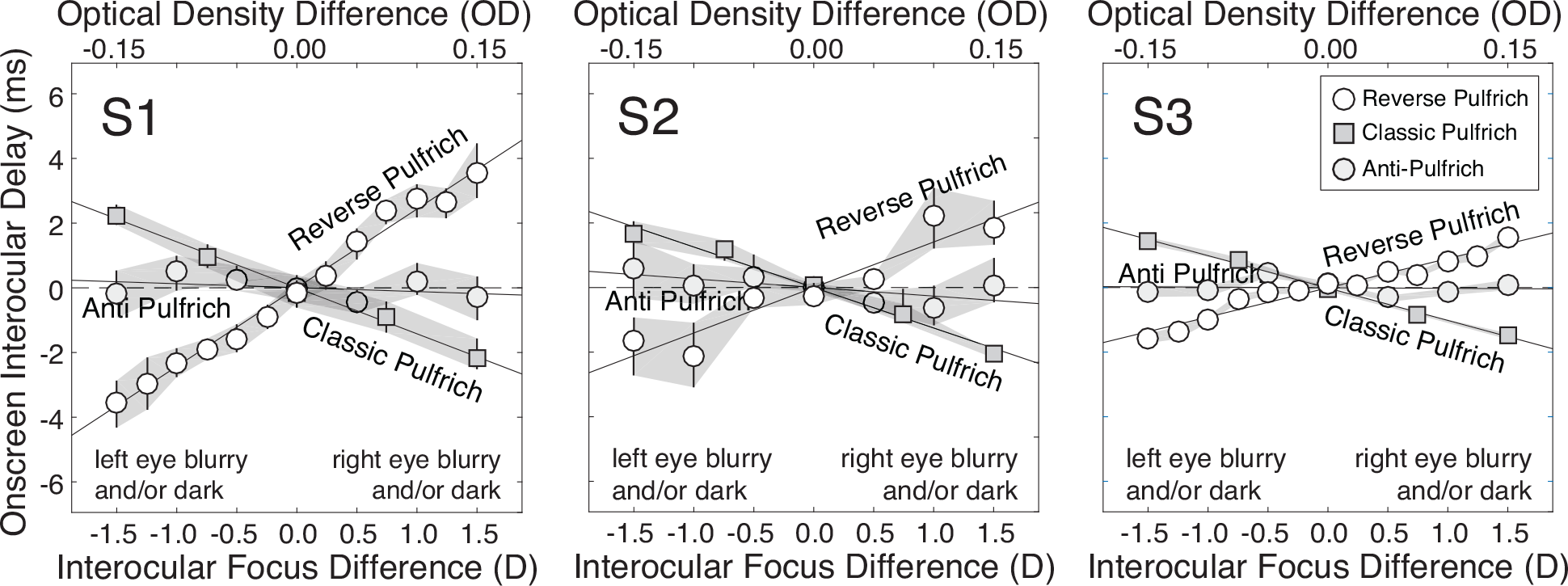
Reverse, classic, and anti-Pulfrich effects for each human observer. Interocular differences in focus error cause the reverse Pulfrich effect; the blurrier image is processed more quickly. Interocular differences in retinal illuminance cause the classic Pulfrich effect; the darker image is processed more slowly. In the anti-Pulfrich condition, the blurry image is darkened to eliminate interocular delay (see Methods).

**Figure S2.**
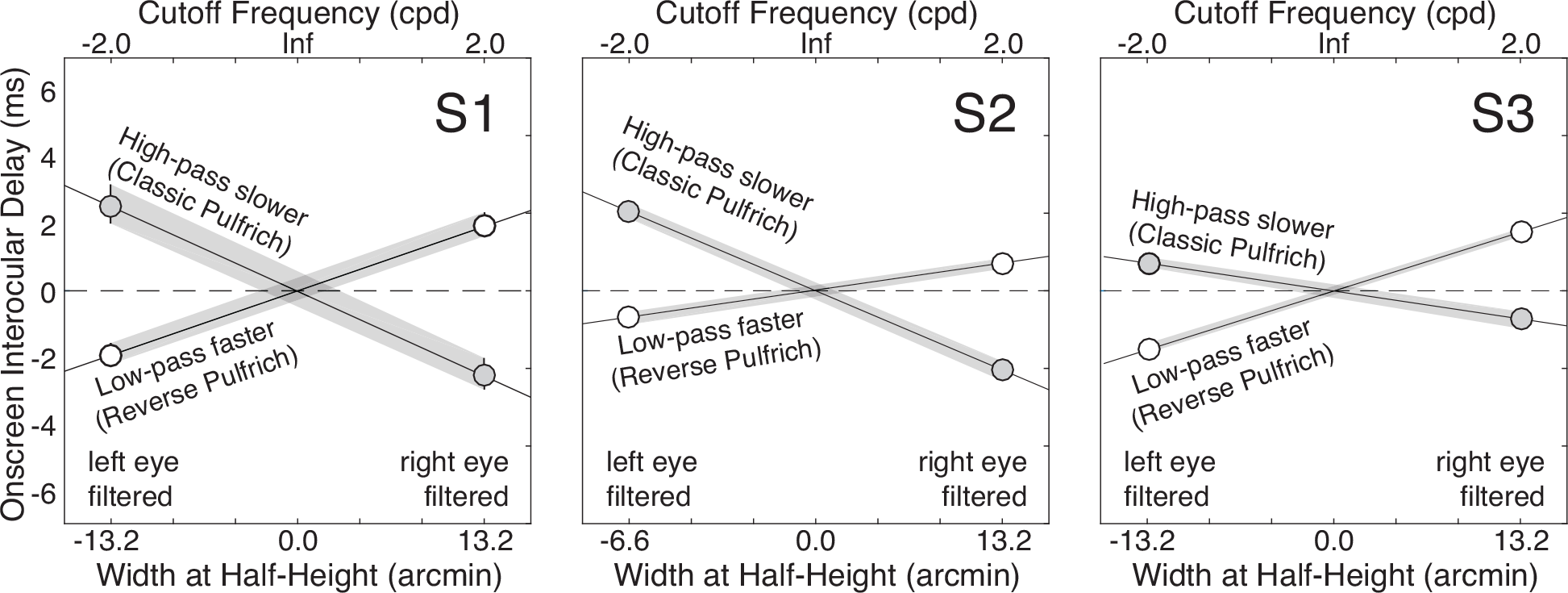
Interocular delays with low-pass filtered and high-pass filtered stimuli for each human observer. The onscreen image for one eye was filtered and the image for the other eye was left unperturbed. Low-pass filtered images were processed faster than unperturbed images, similar to how optical blur induces the reverse Pulfrich effect. High-pass filtered images were processed slower than the unperturbed images. These differential effects cannot be attributed to luminance or contrast because low-pass and high-pass filtered images had identical luminance and contrast (see Fig. S3).

**Figure S3.**
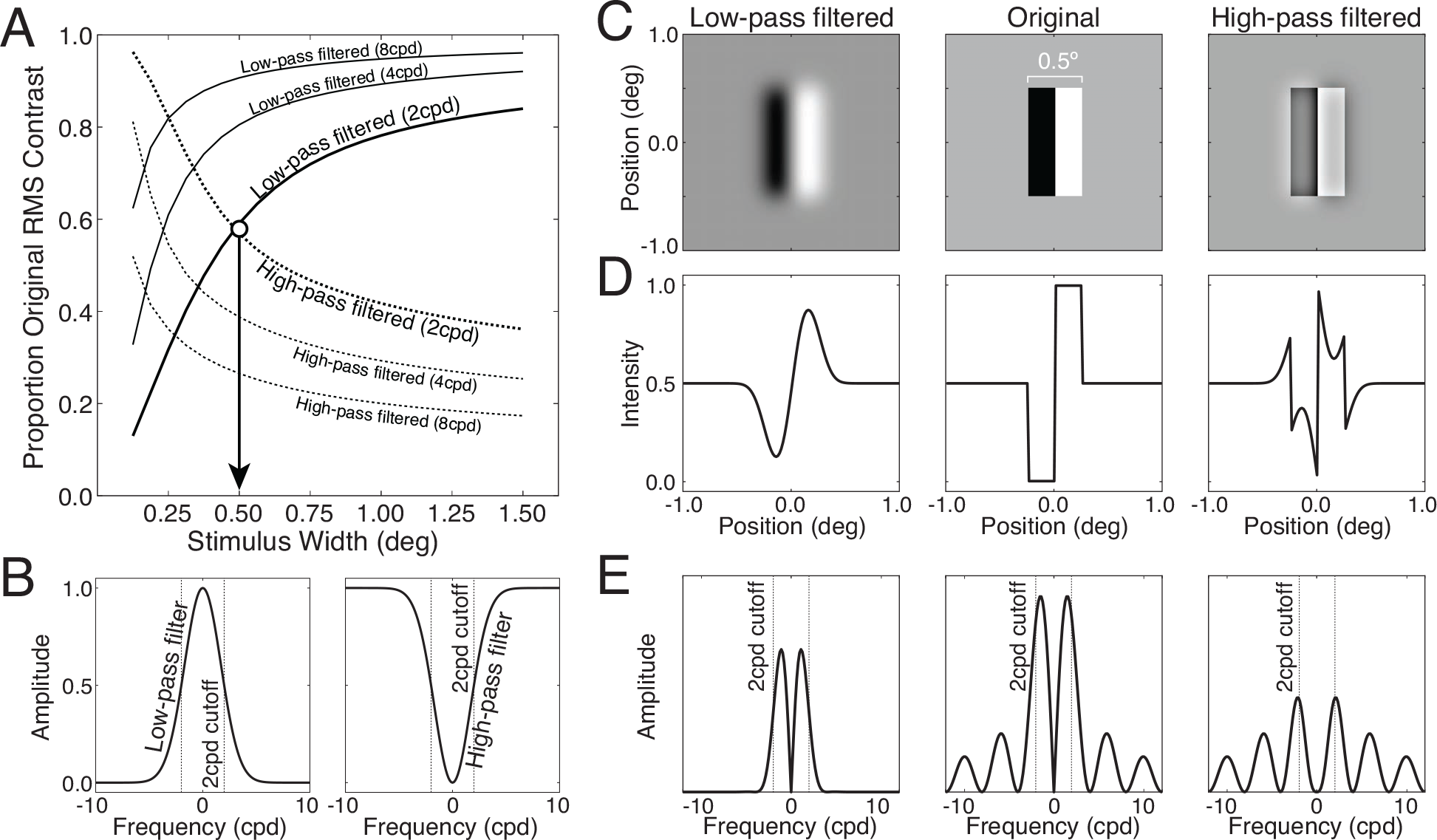
Low- and high-pass filtered stimuli. **A** Proportion of original stimulus contrast after low-pass filtering vs. high-pass filtering (solid vs. dashed curves, respectively) as a function of total bar width. The white circle and arrow indicate the stimulus width (0.5°) that equates the root-mean-squared (RMS) contrast of the stimulus after low- and high-pass filtering. **B** Low-pass and high-pass filters with a 2cpd cutoff frequency. **C** Low-pass filtered stimulus, original stimulus, and high-pass filtered stimulus. **D** Horizontal intensity profiles of the stimuli in C. **E** Amplitude spectra of the horizontal intensity profiles in D. Note how, for each stimulus type, the peak of the lowest frequency lobe shifts relative to the cutoff frequency of the filters.

**Figure S4.**
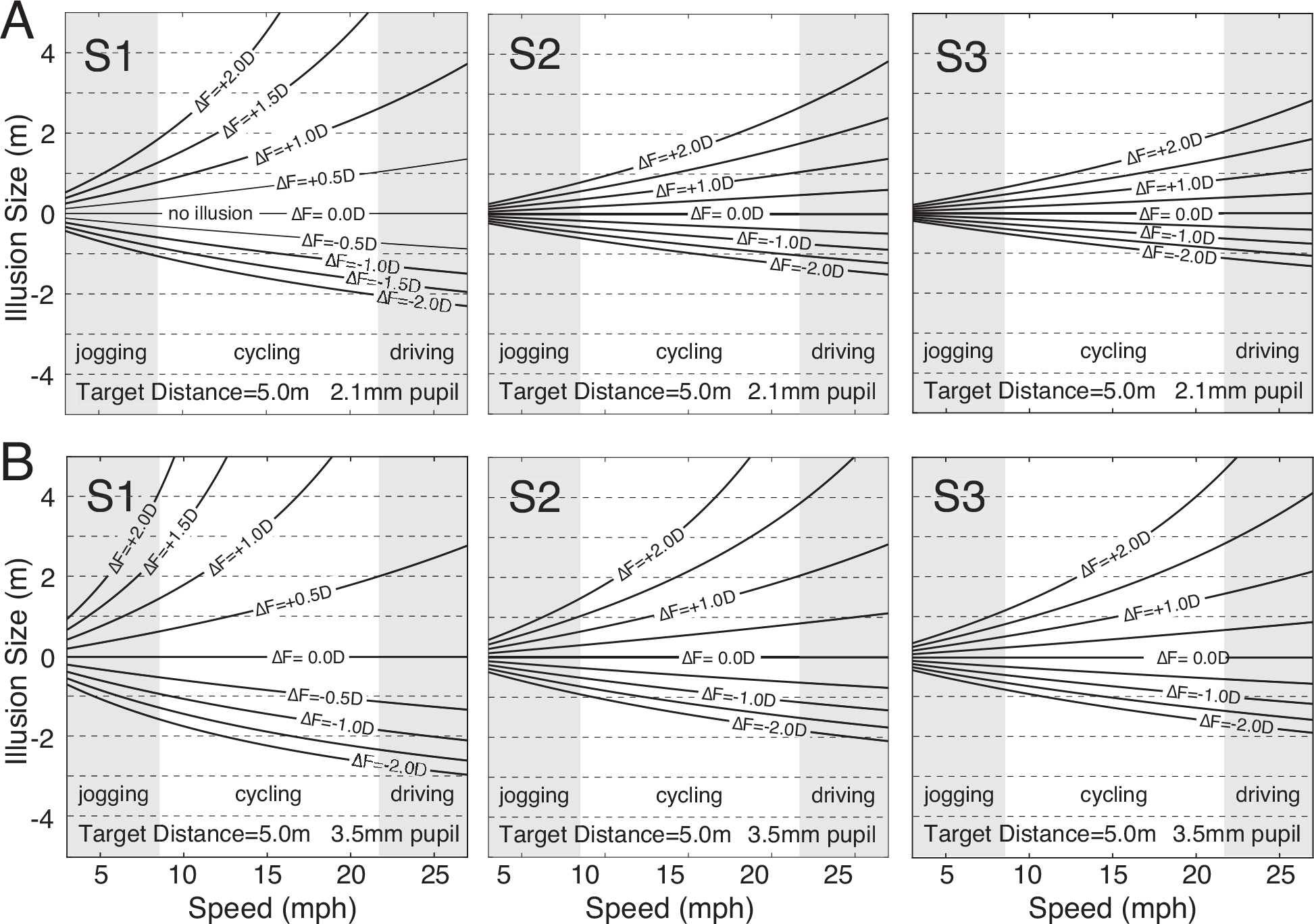
Predicted illusion sizes for different human observers and pupil sizes. **A** Predicted illusion sizes for each human observers under daylight conditions (2.1mm pupil diameter). **B** Predicted illusions for each human observer at dusk (3.5mm pupil diameter). Pupil sizes are consistent with those published by Stockman & Sharpe (2006).

**Figure S5.**
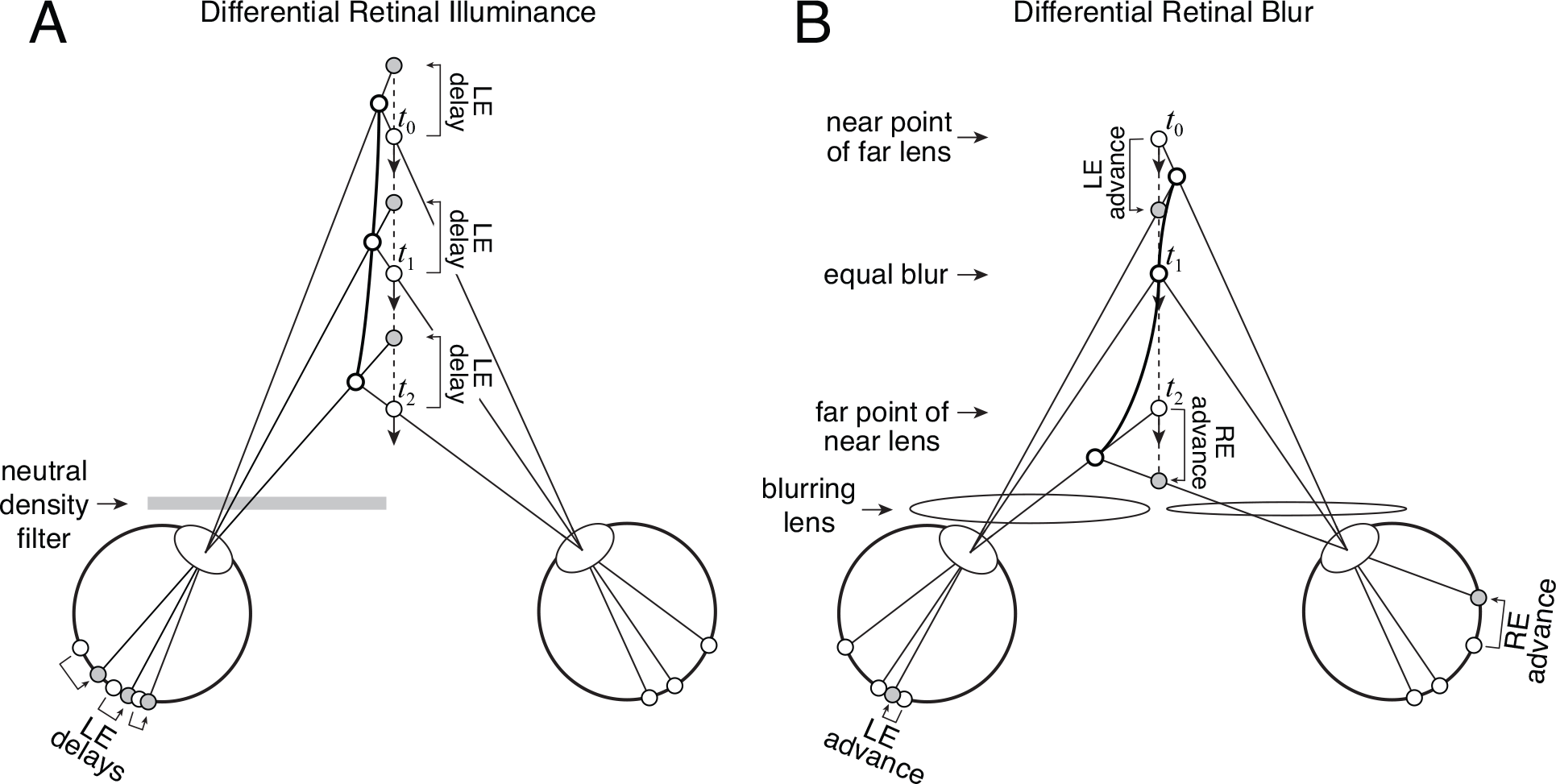
Misperceptions predicted by stereo-geometry for target object motion through depth. **A** Predicted perceived motion trajectory (bold curve), given motion directly towards the observer (dashed line), assuming a neutral density filter causes the left eye image to be processed more slowly. The left eye is processed more slowly independent of the target distance. The illusory motion traces a hyperbolic trajectory^63,69^. **B** Predicted motion illusion, assuming a monovision correction in which the left eye is corrected for near distances and the right eye is corrected for far distances. Here, the eye that is processed more quickly depends on target distance. The illusory motion traces an S-curve trajectory as the target traverses the distances between the near point of the far lens and the far point of the near lens. This occurs because the interocular delay changes systematically as a function of target distance within this range. (Note: the diagrams are not to scale.)

### Misperception of motion towards the observer

Here, we compare the predicted illusory motion trajectories for targets viewed with differential retinal illuminance vs. targets viewed with differential retinal blur. Fig. S5A depicts a target moving directly towards an observer (dashed line) with a neutral density filter in front of the left eye. The target image is processed more slowly in the left eye regardless of target distance. Stereo-geometry predicts that the target will appear to travel along a curved trajectory that bends towards the darkened eye (bold curve) rather than in a straight line^69^. Fig. S5B depicts a target moving towards an observer with a monovision correction where the left eye is focused for near and the right eye is focused for far. Now, the eye that is processed more quickly depends on the target distance. If the object starts at far, the left eye image will be blurry and be processed more quickly. The mismatch in processing speed will make the target appear to bend towards the center of the observer’s head. As the target object arrives at an intermediate distance, where both eyes are equally blurry, the processing speed in both eyes should be identical and the target should appear to be moving directly towards the observer. And as the target object gets to a near distance, the right eye will be blurry and be processed more quickly. Now, the target should appear to be curving to the left. Even more striking effects occur for target objects that are moving towards and to the side of the observer, along oblique motion trajectories. But a full description of these illusions is beyond the scope of the current paper.

**Figure S6.**
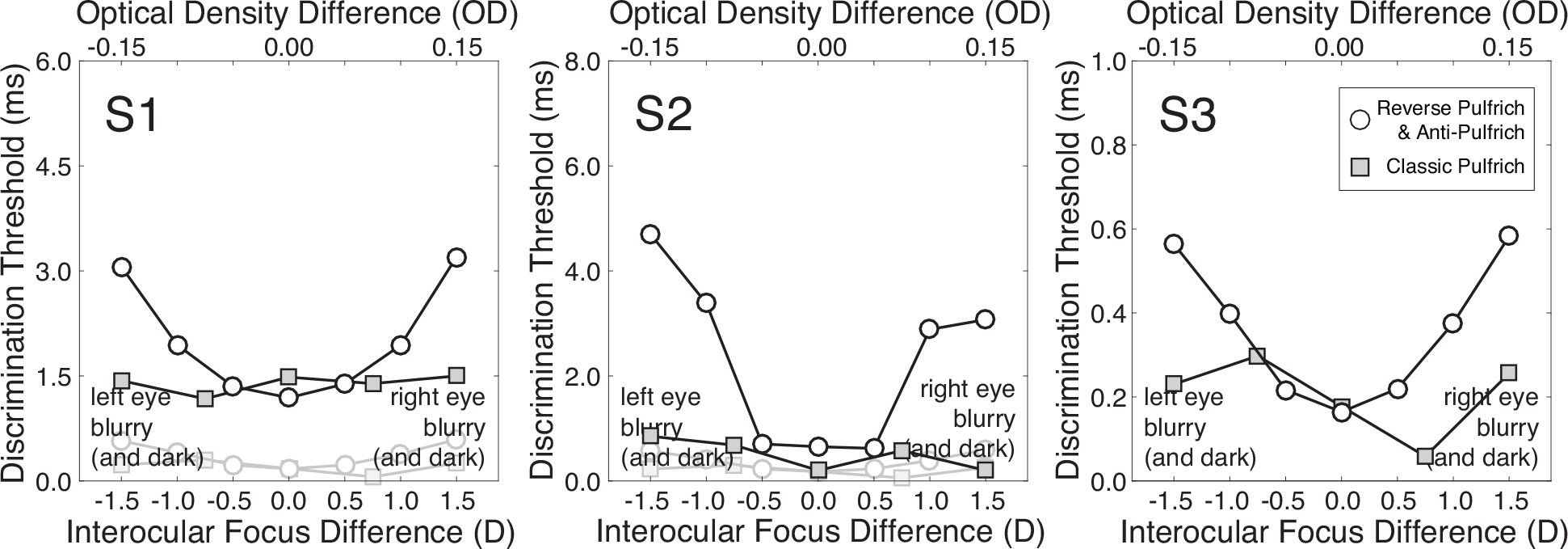
Discrimination thresholds with interocular focus differences. Discrimination thresholds for each observer (*d′* = 1.0) in the reverse Pulfrich conditions (i.e. interocular focus differences only) and the anti-Pulfrich conditions (i.e. interocular focus difference plus interocular retinal illuminance differences) were very similar and were therefore averaged together (white circles). In each human observer, discrimination thresholds increased systematically with differences in interocular blur, consistent with the classic literature on how blur differences deteriorate stereoacuity^63^. These threshold functions thus provide evidence that the desired optical conditions were achieved. Discrimination threshold in the classic Pulfrich conditions (i.e. interocular retinal illuminance differences only) are also shown (gray squares). Differences in retinal illuminance up to +0.15OD had no measurable effect on thresholds. (Note: the y-axis has a different scale for each observer to emphasize the similarities in the threshold patterns. To give a sense of the scale, faint circles and squares in the subplots for observers S1 and S2 replot the classic Pulfrich data from observer S3, the most sensitive observer.)

**Figure S7.**
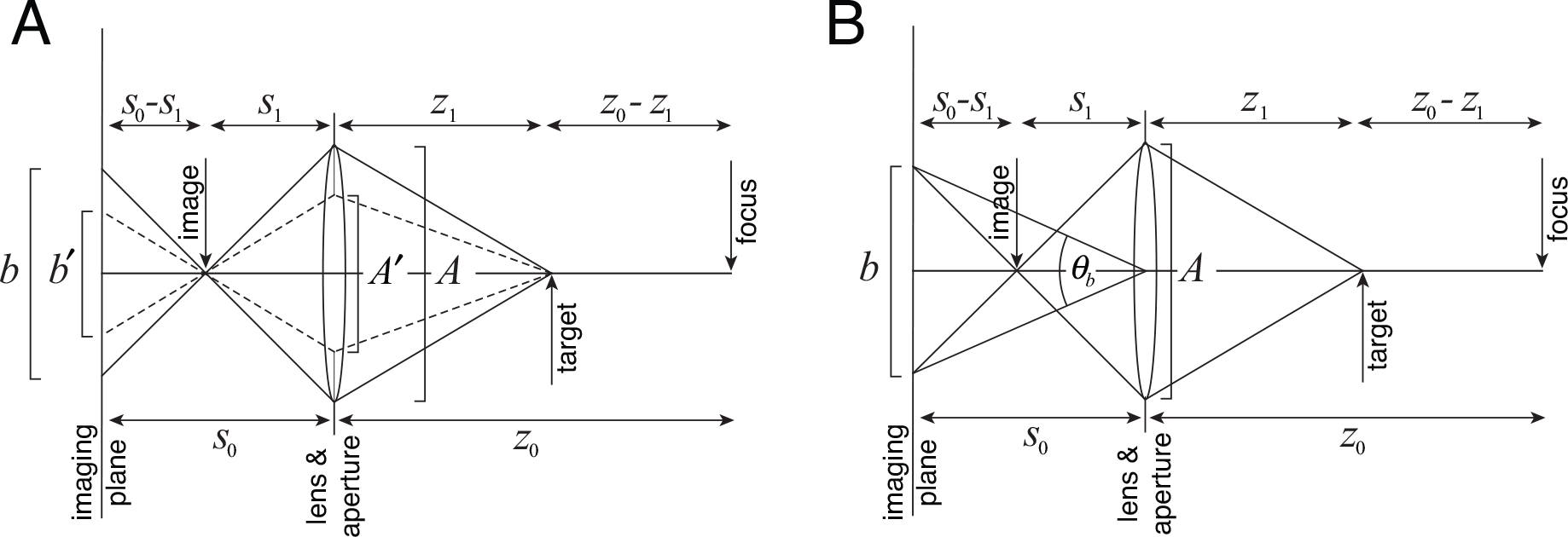
Using geometric optics to relate focus error, aperture size, and blur circle size. **A** Blur circle diameter in meters from aperture and defocus. Solid and dashed lines show how two different aperture sizes (*A* and *A′*) cause two different blur circle sizes (*b* and *b′*) for the same focus error. **B** Blur circle diameter in visual angle. (Note: the diagrams are not to scale.)

### Relationship between focus error and defocus blur in millimeters and in visual angle

Defocus (i.e. focus error) is defined as the difference in dioptric distance between the focus point in object space (i.e. far point conjugated to the imaging plane) and a target

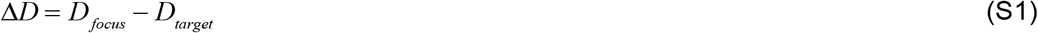

where *D*_*focus*_ and *D*_*target*_ are the dioptric distances to the focus point and to the target. Diopters are defined as inverse meters, so Eq. S1 can be equivalently written

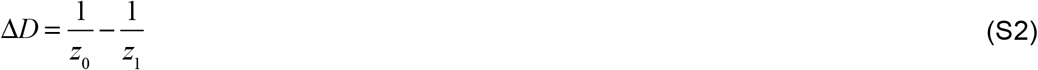

where *z*_0_ and *z*_1_ are distances to the focus point and to the target in meters. The lens equation states that the dioptric difference in object space is equivalently given by the dioptric difference between the imaging plane and the image

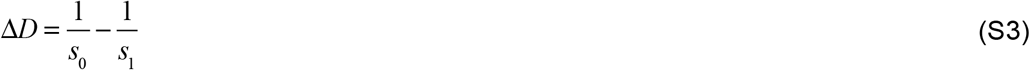

where *s*_0_ and *s*_1_ are distances to the imaging plane and the image in meters. (Note: In this derivation, we use *z* and *s* to be consistent with notational traditions in geometric optics. The symbol *z*_1_ corresponds to the symbol *d* for target distance in the main text.)

Using relations between similar triangles gives the relationship

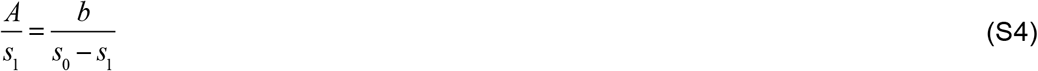

where *A* is the aperture (e.g. pupil) diameter and *b* is the blur circle diameter in meters (Fig. S7A).

Solving Eq. S4 for blur yields

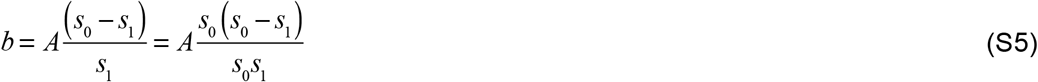

Rearranging Eq. S3 yields

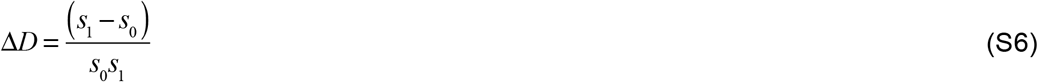

Multiplying Eq. S6 by negative one, taking the absolute value (because the blur circle diameter cannot be negative), and substituting into Eq. S5 gives the blur circle diameter in meters

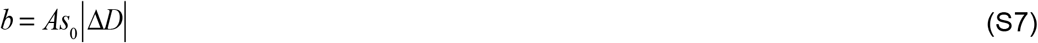

To determine the relationship between the blur circle diameter in meters and in visual angle, it is useful to examine Fig. S7B. From standard trigonometry, the relationship between the blur circle diameter and the subtended visual angle is

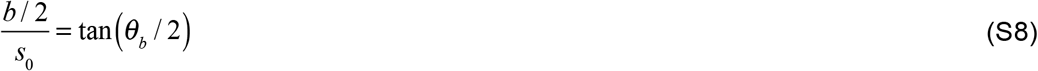

Rearranging after using the small angle approximation

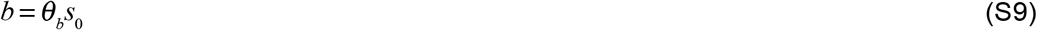

Substituting Eq. 9 into Eq. 7 and solving gives the blur circle in radians of visual angle

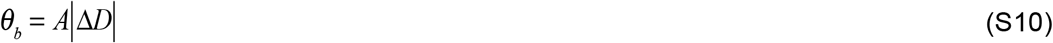

**Figure S8.**
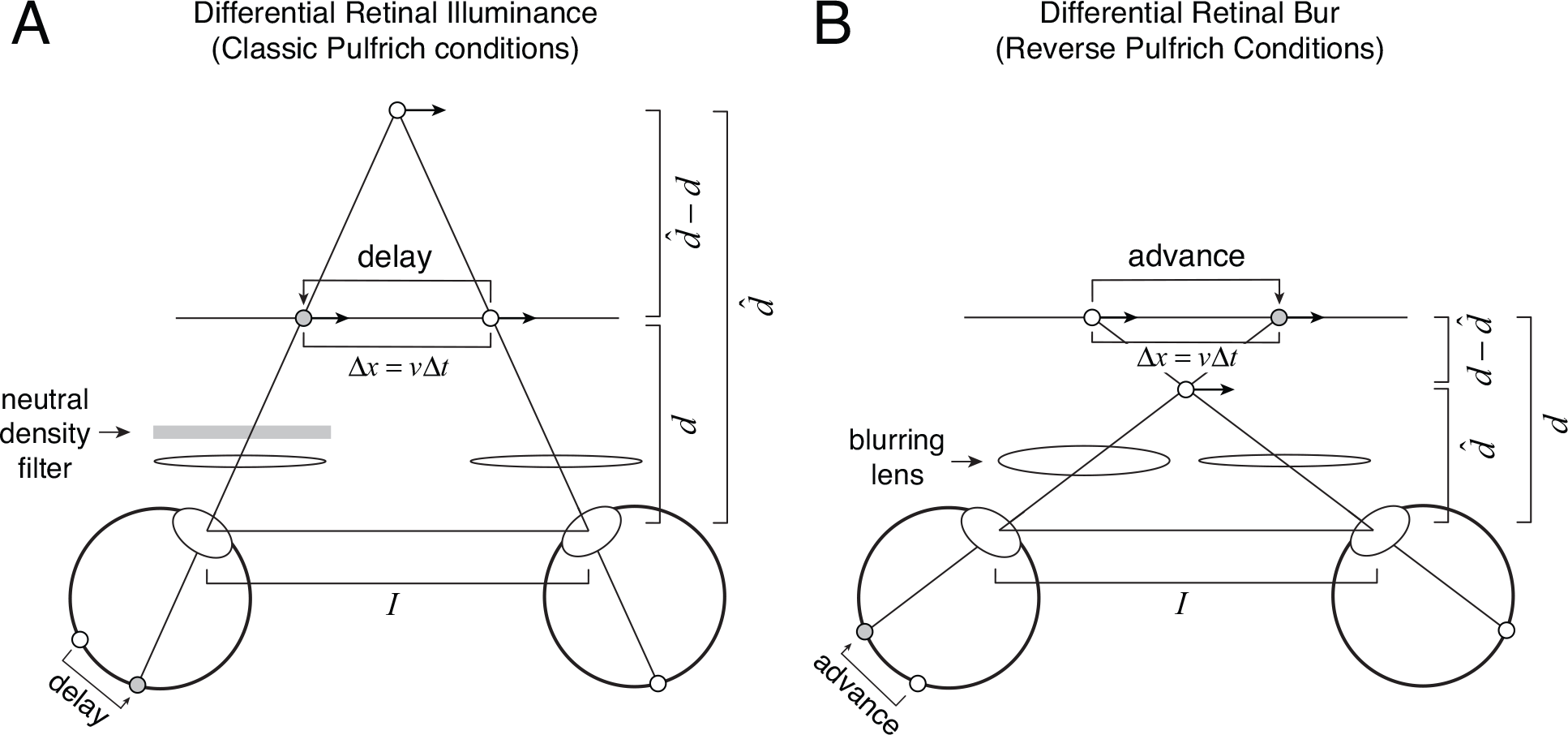
Using stereo-geometry to relate interocular delay, target distance, and illusion size. **A** Stereo-geometry predicting illusion size 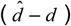 for rightward motion with a neutral density filter in front of the left eye. **B** Stereo-geometry predicting illusion size for rightward motion with a blurring lens in front of the left eye. (Note: the diagrams are not to scale.)

### Relationship between interocular delay, target distance, and illusion size

For a given target velocity and interocular delay the effective spatial offset is given by

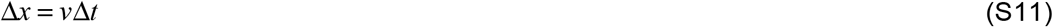

where *v* is target velocity and Δ*t* is interocular delay. The effective spatial offset, target velocity, and interocular delay are all signed quantities. Leftward spatial offsets, leftward velocities, and more slowly processed left-eye images are negative. Rightward spatial offsets, rightward velocities, and more quickly processed left eye images are positive.

By similar triangles, the following relationship holds

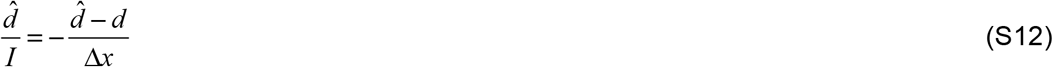

where 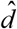 is the estimated (i.e. illusory) target distance, *d* is the actual target distance, and *I* is the interocular distance (Fig. S8AB).

Solving for the illusory target distance yields

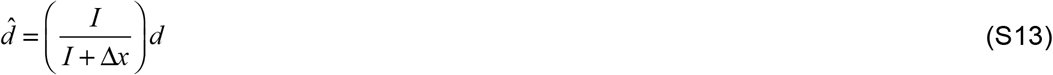

which is identical to Eq. 8 in the main text.

The illusion size 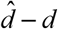 is given by the difference between illusory and actual target distances.

